# Lytic bacteriophages active in urine against multi-drug resistant clinically derived *Klebsiella pneumoniae* causing urinary tract infection

**DOI:** 10.64898/2026.03.23.713486

**Authors:** Ruxandra Calin, Blanca Bernabeu Vilaplana, Judeline Gedeon, Estelle Capton, Cédric Galinat, Azadeh Saffarian, Gautier Pierrat, Yahia Benzerara, Sébastien Wurtzer, Laurent Moulin, Catherine Eckert, Régis Tournebize

**Affiliations:** Mucosal Inflammation and Immunity Team, Université Paris Cité, CNRS, Inserm, Institut Cochin, F-75014 Paris, France & Department of Immunology, Institut Pasteur, F-75015Paris, France; AP-HP Sorbonne Université, Hôpital Tenon, Service Maladies Infectieuses et Tropicales, F-75020 Paris, France; Sorbonne Université, CNRS, Inserm, Centre d’Immunologie et des Maladies Infectieuses, CIMI, F-75013 Paris, France; AP-HP. Sorbonne-Université, Hôpital Saint Antoine, Département Bactériologie, F-75012 Paris, France; AP-HP. Sorbonne-Université, Hôpital Pitié Salpêtrière, Service Bactériologie, F-75013 Paris, France; Eau de Paris, Direction de la Recherche, du Développement et de la Qualité de l’Eau, Eau de Paris, 33 Avenue Jean Jaurès, F-94200, Ivry-Sur-Seine, France; UTechS Photonic BioImaging, Center for Technological Resources and Research, Institut Pasteur, F-75015 Paris, France

## Abstract

**Objectives:** Multidrug-resistant (MDR) *Klebsiella pneumoniae* is an increasingly important cause of recurrent urinary tract infections (UTIs), particularly in high-risk patients such as those with neurogenic bladder, where therapeutic options are limited. Bacteriophage therapy represents a promising alternative, but pre-clinical models and characterization of phages active against UTI-derived strains remain scarce. We therefore aimed to isolate and characterize bacteriophages targeting a clinical MDR *K. pneumoniae* strain causing recurrent UTI and evaluate their activity under urinary conditions.

**Methods:** Three bacteriophages were isolated from environmental samples using an ESBL-producing *K. pneumoniae* clinical isolate obtained from a neurogenic bladder patient. Phages were characterized by genome sequencing, electron microscopy, stability assays, one-step growth curves, and host-range analysis across 79 clinical UTI isolates. Phage activity was quantified in LB medium and human urine using bacterial growth kinetics and a lytic activity score.

**Results:** Three lytic phages from the former siphoviridae family (EDIRA083, EDIRA088, and EDIRA092) belonging to distinct genera were identified. Genomic analysis confirmed the absence of lysogeny-associated, virulence, or antibiotic-resistance genes. Latent periods ranged from 8 to 40 minutes and burst sizes from 38 to 170 virions per infected bacterium. Host-range analysis revealed narrow activity for EDIRA083 and EDIRA088, whereas EDIRA092 infected 29% of the 79 clinical isolates tested. In liquid phage infection assays, overall lytic activity was consistently higher and more sustained in human urine than in LB, suggesting reduced fitness of resistant mutants under urinary conditions.

**Conclusions:** These results identify three genetically distinct lytic phages targeting MDR *K. pneumoniae* and highlight the importance of testing phage activity under infection-relevant conditions. Their activity in urine supports further evaluation of these phages as candidates for therapeutic development against MDR *Klebsiella* UTI.

## Introduction

Each year, millions worldwide face urinary tract infection (UTI), a pervasive bacterial infection that is the second-leading reason for antibiotic prescriptions across the globe [1]. The epidemiological footprint of UTI is substantial. In healthy populations, lifetime risk is 5–15% for men and exceeds 50% for women [2,3]. Approximately 20-50% of patients will experience a recurrent episode within six months [2,4]. Recurrent UTI, defined as at least two episodes in six months or three within one year, are even more prevalent among patient populations with underlying risk factors for UTI and all recurrent UTI patients face increased antibiotic exposure. Risk factors include a variety of pathologies or circumstances ranging from anatomical or functional abnormalities of the urinary tract, indwelling urinary catheters or foreign bodies, to aging, chronic conditions (*e.g.*, poorly controlled diabetes) or immunosuppression [5].

In the general healthy population, uropathogenic *Escherichia coli* is the predominant causative agent of UTI [4]. However, *Klebsiella pneumoniae* (*K. pneumoniae*) accounts for many cases in patients with underlying risk factors [6]. Neurogenic bladder patients, who have bladder dysfunction due to peripheral or central neurological lesions, are among the most vulnerable and have a life-long risk of recurrent UTI, despite optimized management of their bladder dysfunction [7]. In these patients, UTI account for 40% of healthcare-associated infections and are the primary cause of morbidity and the second cause of mortality [8,9]. UTI are also the leading cause for neurogenic bladder patients to seek healthcare and have a major negative impact on quality of life in these patients [10]. Multi-drug resistant (MDR) bacteria cause UTI in up to 50% of these patients, potentially due to repeated antibiotic exposure, with *E coli* and *K. pneumoniae* being the two most prevalent MDR uropathogens [11]. MDR UTI are associated with an increased risk of treatment failure, prolonged hospital stays, higher morbidity and mortality, and a higher risk of renal failure [10–12]. Thus, UTI in patients in this particular risk group are increasing difficult to treat, as the infections no longer respond to standard antibiotic therapy, making them particularly attractive candidates for alternative antimicrobial strategies, such as bacteriophage therapy.

Among MDR uropathogens, *Klebsiella pneumoniae* is of particular concern due to its increasing prevalence and multidrug resistance [13]. The World Health Organization classified MDR *K. pneumoniae* as a critical priority pathogen, highlighting the urgent need for alternative therapeutic strategies as common antibiotics are no longer effective against this pathogen [14]. With recent projections estimating a total of 208 million deaths directly attributable to or associated with MDR organisms between 2025 and 2050 and 10 million annual deaths in 2050, there is an urgent need to develop non-antibiotic based therapies to manage recurrent UTI, especially in the most vulnerable patients [15,16].

Bacteriophage therapy has re-emerged as a viable alternative to conventional antibiotics for treating MDR bacterial infections [17,18]. The advantages of phages as a therapeutic approach include their host specificity, self-replication at infection sites, minimal impact on normal microbiota, and ability to overcome antibiotic resistance mechanisms [18,19]. Phages can be used singly, in cocktails that target different bacterial receptors or strains, or in combination with antibiotics. Several case reports show that phages, alone or in combination with antibiotics can successfully treat recurrent UTI caused by MDR *K. pneumoniae*, however these can be viewed as proof-of-concept, rather than established alternative therapies [20–22]. Moreover, a recent retrospective study of 100 cases of phage therapy in Europe indicated that personalized bacteriophage treatment lead to clinical improvement and bacteria eradication in 77% and 61% of the cases respectively [17]. In contrast, larger studies tend to use a “one size fits all” approach with less convincing efficacy [23]. While “one size fits all” products may comply with stringent good manufacturing practices (GMP), tailored approaches that identify and optimize phage targeting specific clinical strains offer the possibility to adapt or engineer phages to counter resistance emerging during treatment [17,24–28]. Personalised approaches are particularly relevant in UTI due to the diversity of strains that cause infection [29]. Targeted approaches are also needed for *in vivo* modelling to determine how phage and bacteria interact at the site of infection and when exposed to local defense and immune mechanisms specific to each organ. Indeed, models of phage therapy for UTI are limited. In the case of *K. pneumoniae,* while clinical applications for phage therapy target chronic UTI, pre-clinical *in vivo K. pneumoniae* models mainly focus on other types of infection, such as peritonitis, pneumonia, or bacteriemia [30]. Thus, modelling interactions of clinically-derived MDR *K. pneumoniae* causing UTI with therapeutic phages is needed to guide treatment strategies.

In this study, we identified, selected, and characterized new phages that target clinically derived *K. pneumoniae* MDR strains isolated from a recurrent UTI in a neurogenic bladder patient. Because phage candidates are typically evaluated using assays performed in nutrient-rich laboratory media that may not reflect infection-site conditions, we assessed phage activity not only in standard broth but also in human urine. Phages were selected based on their host range, genomic sequence, stability and activity. By comparing the activity of phages combined into a therapeutic cocktail in laboratory media and urine, we determined their therapeutic potential for future *in vivo* evaluation.

## Methods

### Bacterial strains

Strain RT535 is a *K. pneumoniae* isolate from a neurobladder patient under care at the Assistance Publique – Hôpitaux de Paris (APHP) - Hospital Tenon Paris, France that was used to isolate bacteriophages. All other *K. pneumoniae* strains are strains isolated from patients with UTI at AP-HP hospitals (Paris, France). Strains were collected in the framework of an epidemiology study KPCartophage (APHP231314). All strains were grown in LB-Lennox broth or agar (Invitrogen).

### Bacteriophage isolation, amplification and titration

Bacteriophages EDIRA083, EDIRA088 and EDIRA092 were isolated from sewage water from Saint-Thibault-des-Vignes, France (Provided by Eau de Paris); the Seine River, sampled in Paris, France; and the Auvezère River, sampled in Cherveix-Cubas, Dordogne, France, respectively. Phages were directly isolated from sewage water concentrated by centrifugation though sucrose cushion gradient (EDIRA083) or after amplification by growing bacteria in 1x LB-Lennox broth spiked with the river water samples (EDIRA088, EDIRA092). Clear plaques were successively isolated 3 times to obtain pure phages. Phages are stored at 4°C. To amplify phages, *K. pneumoniae* isolate RT535 was grown for 2 hours at 37°C, 180 rpm agitation to reach exponential growth phase. Bacteria were diluted to OD=0.2, phages were added, and the culture grown overnight at 37°C shaking. The next day, bacteria were removed by centrifugation, 10 minutes, 4000g, and the bacteriophage-containing supernatant filtered through 0.22 µm filters. Bacteriophages were titrated using a spot assay method. Briefly, bacterial culture at OD between 0.15 and 0.2 were spread onto LB-Lennox agar containing 1 mM CaCl_2_ and 10 mM MgSO_4_. Once the plate dried, serial 10-fold dilutions were spotted onto bacteria lawns, the plates incubated overnight at 37°C and the number of plaques enumerated. Efficiency of plating were determined as the ratio of titer of phages measured onto test strains relative to the titer measured with the isolation host strain. Ratio are expressed as Log10 ratio.

### Genome sequencing and annotation

Bacteria and phage genomes were sequenced using Illumina technology at the AP-HP GenoBiomics core facility (Créteil, France). Briefly, for phage sequencing, filtered bacteria lysates containing phages (titer > 10^9^ PFU/mL) were diluted in 10 mM Tris-HCl pH7.5, 2.5 mM MgCl_2_, 0.1 mM CaCl_2_ containing 50 µg/mL DNAse I (Merck) and incubated at 37°C for an hour to digest contaminating bacterial DNA before addition of 4 mM EDTA and 0.08 % SDS. Phages or bacteria suspensions were then processed for DNA extraction and Illumina sequencing at the GenoBiomics platform (paired end, 150 bp).

For genome assembly, read quality was assessed using fastQC [31], trimming performed using Trimmomatic [32], and assembly using Spades [33]. A prior down-sampling of raw reads using seqtk [34] was required for the assembly of the phage genomes. Bacterial genomes were annotated using Bakta [35], plasmids identified using PlasmidFinder [36], capsule and LPS type and antimicrobial and virulence factor presence determined by Kleborate [37,38], and presence of bacterial defense systems assessed by DefenseFinder [39–41]. Phages genomes were annotated with Pharokka [42] and phold [43], therapeutic suitability assessed using PhageLeads [44]. Phages diversity was assessed using VipTree [45] and taxonomy determined using taxMyPhage [46].

### Electron microscopy

500 µl of phages with titers higher than 10^9^ PFU/mL were centrifuged 75 minutes at 20,000g at 4°C, resuspended in 10mM Tris HCl, pH7.5, 1mM CaCl_2_, 10mM MgSO_4_ (SM buffer) and spun again for 30 minutes at 20,000g at 4°C. Phages were resuspended in 50 µl SM buffer and kept at 4°C. Phages were spotted on an EM carbon grid, stained with 2% uranyl acetate and observed using a Tecnai F20 microscope. Imaging was done at the Ultrastructural BioImaging core facility at Institut Pasteur.

### pH and temperature stability

To test pH stability, phage suspensions were diluted 1/10 in LB Lennox buffered at pH3, pH5, pH7, pH9 or pH11 and incubated overnight at 37°C prior to titration. Temperature stability was assessed after incubating phage suspensions at 37, 45, 50, 55, 60, 65, or 70°C for 1 hour. Phages were titrated by spotting dilutions onto a lawn of RT535 bacteria as described above.

### One-step growth curve

Mid-log phase culture of host bacteria was diluted to OD=0.25 and infected with phages at a Multiplicity of Infection (MOI) 0.1. After 5 minutes of co-incubation, the culture was diluted 1/1000 to prevent subsequent reinfection. Aliquots from the culture were retrieved and kept on ice at intervals of 2 minutes until 15 minutes, then every 5 minutes until 60 or 90 minutes post-infection. Each aliquot was titrated to determine the number of PFU/mL over time. Burst size, the expected number of virions produced by one infected cell over its life-time, and latent period, the time a phage particle needs to reproduce inside an infected host cell, were measured for each phage.

### Lytic Activity Score

Cultures of *K. pneumoniae* isolate RT535 at OD=0.2 were infected with phages at an MOI of 0.1 in 160 µl volume in 96 well plates and incubated at 37°C for 46 hours with the OD600 measured every 15 minutes using a heated Infinite M200 spectrophotometer (Tecan). Bacteria and phages were grown in LB-Lennox broth or pure urine (urine pooled from 3 female donors aged under 25 years, with no leukocyturia, not taking contraceptive and more than 1 week apart from their period or a pool from 1 female and 1 male healthy donors). All healthy donors did not take any antibiotics at least one month before. Growth curves were used to determine the area under the curve (AUC) using Prism 10 (Graphpad Software) and to calculate the Lytic Activity Score as (AUC _bacteria_ -AUC _bacteria+phage_) / AUC _bacteria_ x100.

## Results

### Three environmentally-sourced phages target clinical *K. pneumoniae* strains

To identify phages targeting *K. pneumoniae*, we first selected a clinical strain isolated from a neurobladder patient suffering from recurrent UTI. The strain, RT535, is an Extended Spectrum Beta-Lactamase (ESBL)-producing isolate belonging to Sequence Type (ST) ST1958, carrying the capsule locus KL110, LPS serotype O2a, and without virulence-associated factors, such as the siderophores aerobactin, salmochelin and yersiniabactin, the colibactin toxin and regulators of expression of capsule. It carries both IncFIB(K) and IncFII(K) type plasmids and expresses a CTX-M-15 beta-lactamase. The strain is resistant to fluroquinolones, aminosides, trimethoprim/sulfamethoxazole, and fosfomycin. RT535 carries genes encoding for two type_II, one type_IIG_2 and a type_IV restriction modification defense systems, and one of each of the PD-T4_3, Hachiman, Cas, CBASS, Septu, AbiE, DS-20 and Tg GGvAB systems [47]. No intact prophages were identified in the genome.

Three phages, named EDIRA083, EDIRA088, EDIRA092, were isolated from sewage and environmental samples (**Figure 1**). After 16 hours of culture, phages EDIRA083 and EDIRA088 developed small plaques on bacterial lawns of strain RT535 of 0.24±0.03 and 0.19±0.02 mm diameter respectively, while phage EDIRA092 had slightly larger plaques of 0.38±0.09 mm. A slow depolymerase activity was observed with EDIRA083 but depolymerase halo starting to be visible from 2 days of incubation of the plate, indicating that this phage encodes for an active KL110 capsule-specific depolymerase. No depolymerase halo was observed for EDIRA088 and EDIRA092.

**Figure 1.**
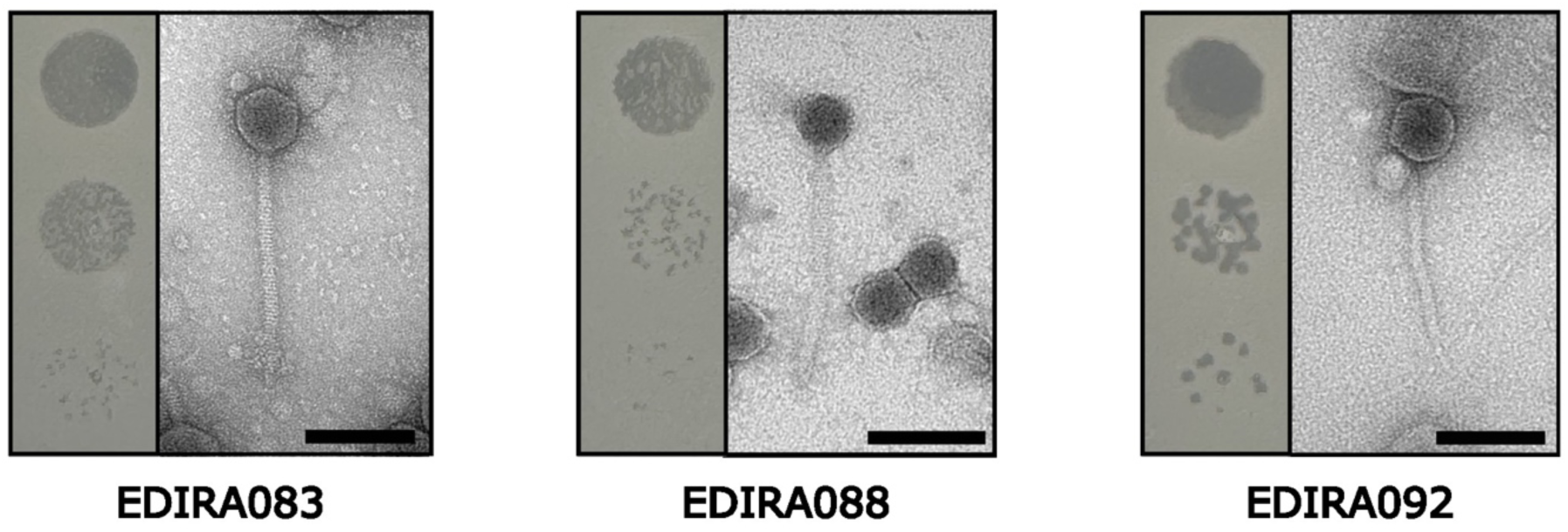
Three phages isolated from environmental sources have activity against a clinical *K. pneumoniae* isolate. Representative plaque morphology of phages EDIRA083, EDIRA088 and EDIRA092 spotted on bacterial lawns of *K. pneumoniae* clinical isolate RT535 (left side of photos) and electron microscopy images of each phage. Scale bar: 100 nm.

We observed bacteriophage morphology by electron microscopy. All three phages were of the siphoviridae morphotype with similar sized capsids, ranging from 54.9 nm to 58.8 nm by and 51.9 nm to 54.2 nm (**Table1** and **Figure 1**). The tail of EDIRA088 was notably longer (219.1 nm) and wider (17.8 nm) than those of EDIRA083 and EDIRA092 (**Table1** and **Figure 1**).

#### EDIRA083, EDIRA088, and EDIRA092 phages are genetically distinct

We next sequenced the genomes of the three phages isolated. Genome sequencing of EDIRA083, EDIRA088, and EDIRA092 showed genome sizes of 57.8 kb, 48.1 kb, and 50.6 kb, encoding for 88, 63 and 91 proteins, respectively (**Figure 2A**). Predicted hypothetical proteins represented 53%, 50%, and 52% of the annotated proteins in EDIRA083, EDIRA088 and EDIRA092, respectively. No lysogenic-related genes, such as transposases or integrases, or antibiotic-resistance or bacterial virulence genes were identified, probing ABRicate database, indicating that these phages may be suitable for therapeutic application and will not be excluded for their recombination or antimicrobial resistance transmission potential. Whole genome proteomic clustering using VipTree showed that the three phages were phylogenetically distinct, belonging to different genera (**Figure 2B**). Classification of the bacteriophages using Taxmyphage indicated that EDIRA083 and EDIRA092 were part of the Yonseivirus and Peekayseptimavirus genera respectively, whereas BLAST analysis showed that EDIRA088 was 97.8% identical with 95% coverage to the vb_Kpn_K34PH154 bacteriophage, a member of a genus yet not taxonomically defined within the Caudoviricetes class.

**Figure 2.**
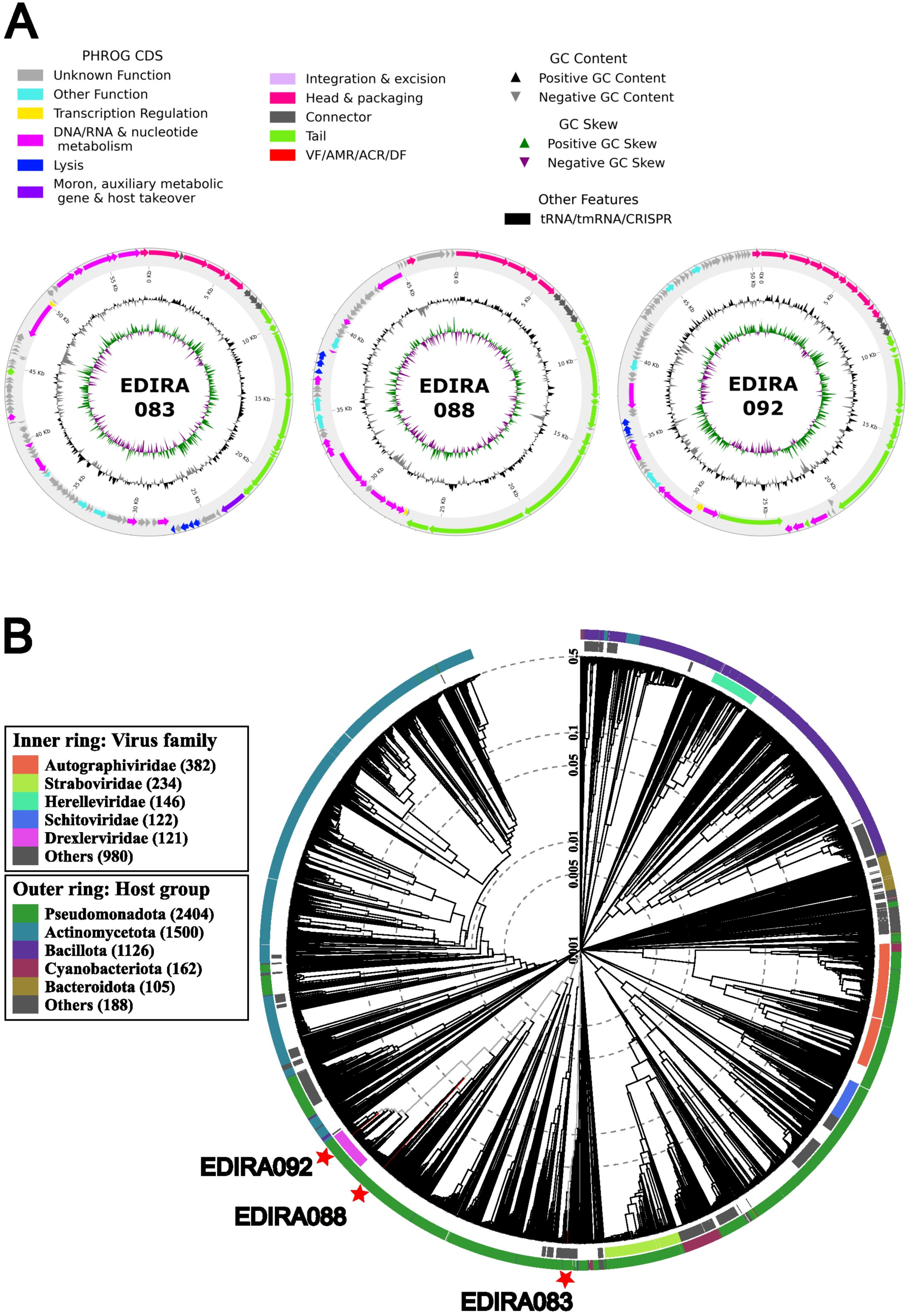
EDIRA083, EDIRA088, and EDIRA092 are genetically diverse. A) Genomic maps and functional annotation of phages EDIRA083, EDIRA088 and EDIRA092 were made using pharokka followed by phold, ORFs are colored according to PHROG annotation. B) The proteomic tree was generated using VipTree, and shows EDIRA083, EDIRA088, and EDIRA092 clustering among reference phage database.

#### EDIRA083, EDIRA088, and EDIRA092 phages are stable at physiological pH and temperature

To characterize infection dynamics of the bacteriophages, we used one-step growth curves. Phages EDIRA083 and EDIRA088 had a latent period of 40 and 25 minutes and a burst size of 170 and 43 virion particles produced per bacterium, respectively. The Peekayseptimavirus EDIRA092 phage had a very short latent period of 8 minutes and a burst size of 38 virion particles produced per bacterium (**Figure 3A**). All three phages were completely inactive at pH3, stable at pH values between 5 and 9, and partially inactive at pH11, showing that they are stable at the pH values of most sites of host infection, including bladder and urine. All phages were stable up to 55°C. EDIRA088 was completely inactivated at 60°C, while EDIRA083 and EDIRA092 were still stable at 60°C. EDIRA083 and EDIRA092 were completely inactive at 70°C.

**Figure 3.**
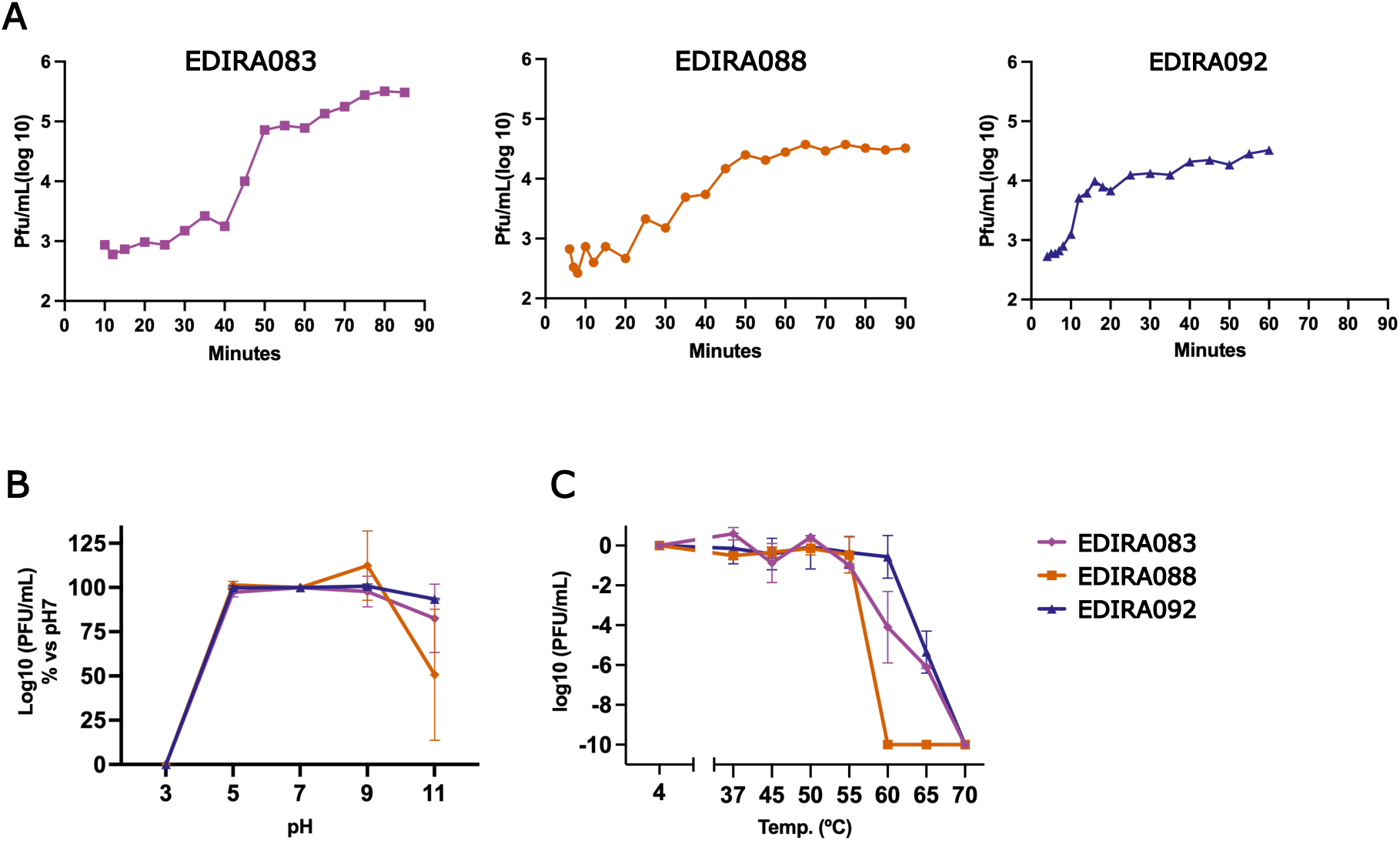
EDIRA083, EDIRA088 and EDIRA092 are active at physiological pH and temperature. A) Representative one-step growth curves of EDIRA083, EDIRA088, and EDIRA092. Each phage was co-incubated with strain RT535 for phage adsorption, and then diluted to limit phage-bacteria interaction and samples were collected at regular intervals to determine the latent period and burst size. Data are representative of 2 independent experiments. B) pH stability was measured after incubating phages over a range of pH for 16 hours and determining phage titer relative to titer after incubation at pH7. Data are the mean of 2 to 3 experiments. C) Temperature stability was measured after incubation of phages over indicated temperatures for 1 hour followed by phage titering. Data are normalised to values obtained at 37°C and expressed in log10 reduction relative to 37°C values. Data are the mean of 3 experiments.

### Host range of EDIRA083, EDIRA088, and EDIRA092

We next assessed the host range of the isolated phages by determining their efficiency of plating, *i.e.* their capacity to infect and produce a reproductive infection onto a specific strain relative to the parental strain, towards a collection of 79 *K. pneumoniae* strains isolated from the urine of patients treated at Paris area hospitals. These clinical strains represented 57 different STs and 43 different capsule types (**Figure 4**). We observed that EDIRA083 and EDIRA088 phages had an extremely narrow host range, in which EDIRA083 was lytic on 2 clinical *K. pneumoniae* strains and EDIRA088 targeted only the RT535 strain used to isolate this phage (**Figure 4**). Conversely, the phage EDIRA092 recognised 29% of strains, with an efficiency of plating relative to RT535 ranging from -3.5 log10 to 1 (**Figure 4**). Phage activity was not correlated with specific capsule or O-antigen type or the ST (**Figure 4**).

**Figure 4.**
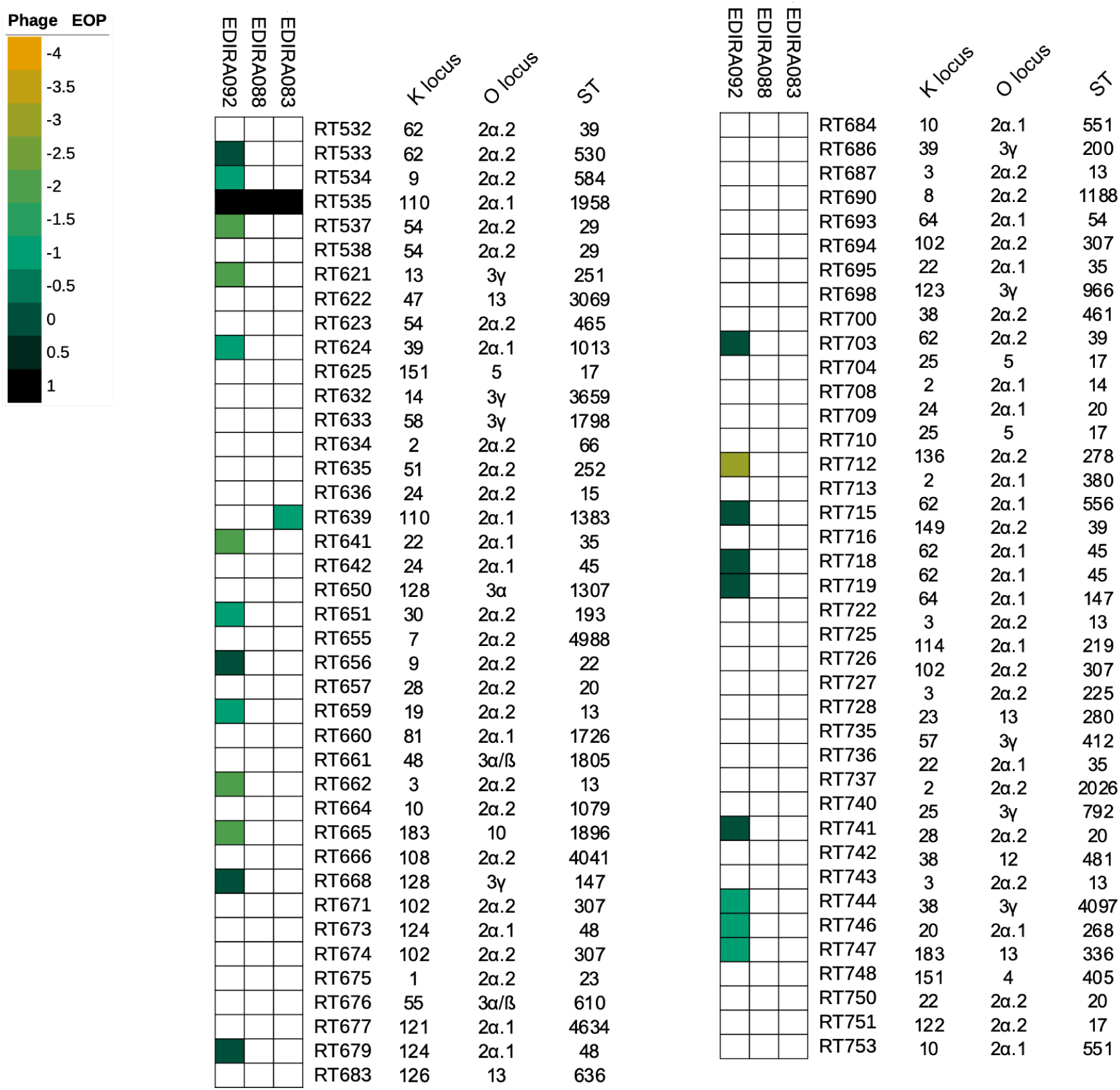
EDIRA083, EDIRA088 and EDIRA092 have distinct host ranges. The activity of phages EDIRA083, EDIRA088 and EDIRA092 was evaluated on a collection of *K. pneumoniae* strains isolated from patients with UTI. Activity was quantified by measuring the efficiency of plating. Data are expressed by color scale as log10 reduction compared to the activity on strain RT535. ST, KL- and O-loci types are indicated for each strain.

### Phage EDIRA088 targets *K. pneumoniae* better than other phages in urine

As the activity of phages can be modified in biological tissues, we investigated the activity of the three phages in LB and human urine by measuring the growth kinetics of RT535 in these media and determining a Lytic Activity Score (LAS). This score quantifies phage lytic activity, with a high score indicating a high lytic activity and stronger suppression of bacterial growth. *In vitro* in LB, we observed that the three phages, singly or in combination, reduced bacterial growth in the first 8-10 hours before observing regrowth of the phage-resistant bacteria up to 46 hours (**Figure 5A**). As growth kinetics are often performed only over 24 hours, to compare our results with previous published data, we quantified the LAS at both 24 and 46 hours of culture (**Figure 5B**). In three independent experiments, we observed that the LAS of EDIRA083 and EDIRA088 were similar at 24 hours with a LAS of around 66-70%, while the values were more variable (59 - 79%) for EDIRA092, reflecting of the stochastic emergence of phage-resistant bacteria in these experiments. This difference in LAS values across experiments was also observed with higher variability when phage EDIRA092 was used in 2- or 3-phage cocktails (**Figure 5B**), indicating that EDIRA092-resistant bacteria can emerge quickly when they are grown in LB. Of note, in some experiments, no phage-resistant bacteria were observed, with LAS values measured between 92 and 96%. In addition, as phage-resistant bacteria grew efficiently once they emerged, LAS values were globally lower after 46 hours of cultures (**Figure 5B**).

**Figure 5.**
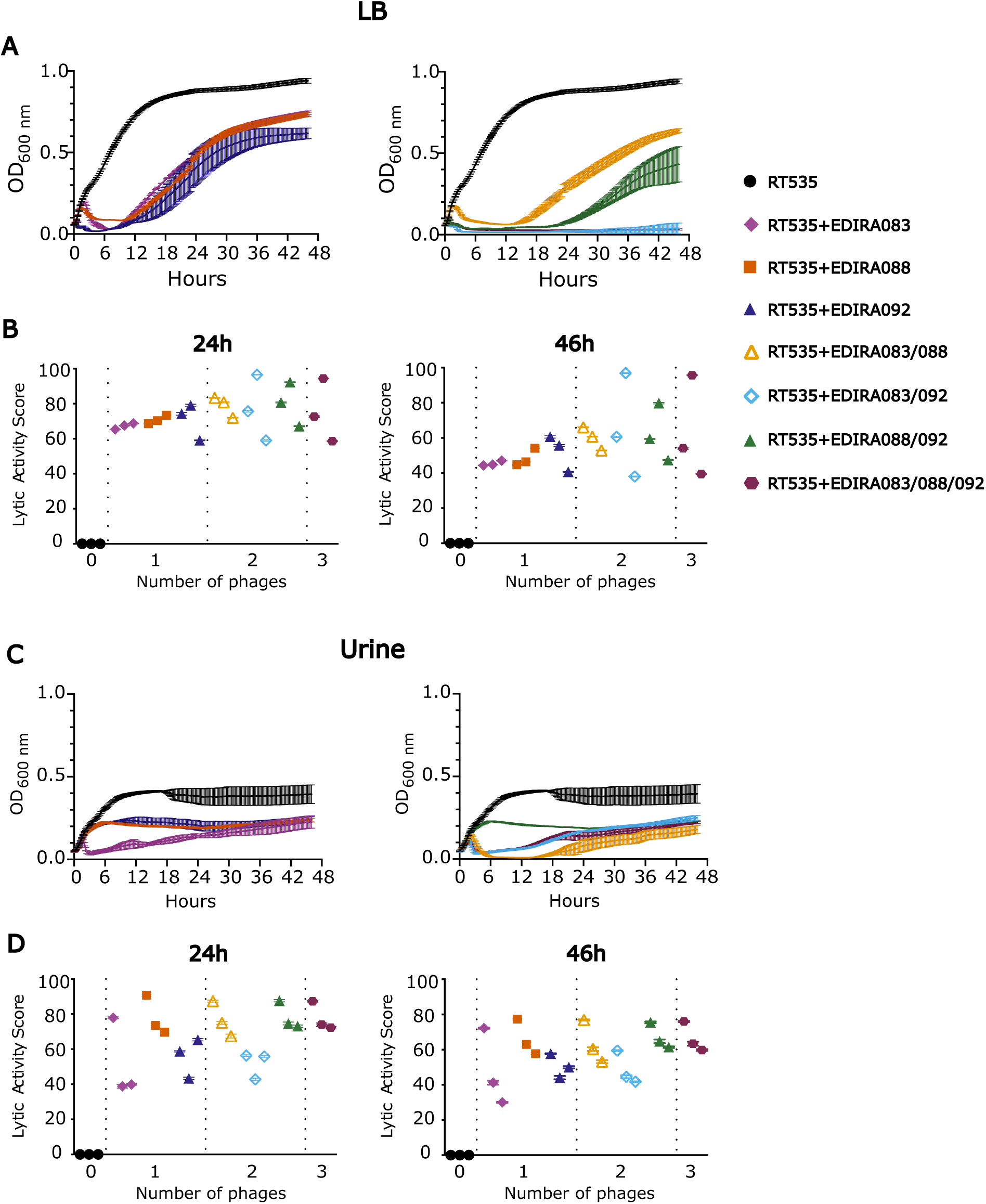
EDIRA088 outperforms other phages in urine. Lytic activity of phages in liquid cultures was determined by growing *K. pneumoniae* strain RT535 in LB (A, B) or urine (C, D). Representative growth curves show RT535 strain in the presence of single phages EDIRA083, EDIRA088, EDIRA092, or in combination in LB or urine over 46 hours growth. The Lytic Activity Score (LAS) was determined for each of 3 independent experiments performed in LB or urine after 24 or 46 hours. High values indicate high lytic activity and low growth of phage-resistant bacteria. Data plotted are average ± sd of 3 technical replicates for each of 3 independent experiments.

Bacteria growth was more limited in urine compared to LB, however emergence of phage-resistant bacteria was also observed (**Figure 5C**). The LAS of single phages was more variable than in LB after 24 and 46 hours of infection and was globally higher at 46 hours in urine than in LB. Interestingly, phage EDIRA088 outperformed EDIRA083 and EDIRA092 in urine when used singly (**Figure 5D**). Cocktails that included EDIRA088 had higher LAS compared to cocktails that did not include this phage (**Figure 5D**). Overall, the activity of the phages was higher in urine than in LB, suggesting that the fitness of the phage-resistant bacteria is lower in urine, thus supporting of the potential of these phages for treatment of UTI.

## Discussion

As the second cause of antibiotic prescription, UTI contribute to driving antibiotic resistance. Indeed, in patients with complicated recurrent UTI caused by MDR bacteria, such as in neurogenic bladder patients, treatment failure is associated with repeated hospitalization, the use of reserve antibiotics, and a therapeutical impasse [48]. Such patients need new therapeutic alternatives. Phage therapy has therapeutic potential, but *in vivo* models are rare and studies in humans are predominantly case reports [17,18,49,50]. To identify phages active against UTI-relevant bacteria, we characterized three lytic phages active against an ESBL-producing *K. pneumoniae* strain from a patient with neurogenic bladder and recurrent UTI. We isolated 3 phages belonging to different genera from sewage and environmental samples. Their host range profiles and lytic kinetics suggest these phages have different mechanisms of action in targeting *Klebsiella*.

The three phages described here are identified as *Yonseivirus* (EDIRA083), *Peekayseptimavirus* (EDIRAO92) and belonging to a genus not yet taxonomically defined (EDIRA088). EDIRA088 is 97.8% identical with 95% coverage to the phage vb_Kpn_K34PH164. All three phages had no transposases, integrases, or antibiotic-resistance or bacterial virulence genes. They all exhibited pH and temperature stability within physiological ranges. These features support the potential use of these phages in therapeutic applications, making them theoretically suitable for the treatment of UTI. Additionally, burst size and latent period both influence therapeutic success. *In vitro* studies on biofilm show that phages with higher burst sizes and shorter latent periods better control biofilm formation, which may be relevant in UTI [51]. Our phages showed latent periods ranging from 8-40 minutes with burst sizes from 38 - 170 virions per bacterium. A short latent period, even with a small burst size, such as that seen in phage EDIRA092, might be advantageous for the treatment of an active infection with high bacterial colonization. However, in the context of low host bacteria density, longer latent periods and larger burst sizes (as for phage EDIRA083) could be more favourable [52].

A narrow host range is one of the main challenges when searching for phages against UTI pathogens, such as *E coli* or *K. pneumoniae*. Indeed, as opposed to other bacteria, such as *Staphylococcus aureus*, *K pneumoniae* phages generally have narrow host ranges because of the high surface antigenic heterogeneity of *Klebsiella* species, with more than 163 different *Klebsiella* K-types [30,38,53]. Capsule polysaccharide diversity is considered one of the most important determinants of phage host specificity, as the initial phage-receptor interaction involves recognizing the bacterial capsule [53]. Indeed, infection by *Yonseiviruses* PIN1 and PIN2 are dependent on *K. pneumoniae* capsule and O-antigen [54]. Receptors for EDIRA083 and EDIRA092 are not yet known. When tested on 79 *K. pneumoniae* UTI strains from 57 different STs and expressing 43 different capsule types, EDIRA083 and EDIRA088 had an extremely narrow host range. However, EDIRA092, targeted nearly 30% of the tested strains, indicating a comparatively broad host range for a *Klebsiella* phage. Interestingly phages showed different susceptibility for strains with the same K locus and ST, showing the complexity of matching *Klebsiella* phages to susceptible *Klebsiella* strains. However, the host range of EDIRA083 and EDIRA088 could be specifically optimized for specific strains using directed evolution [55–57]. By combining and, potentially, evolving phages with different host range, it may be possible to extend the host range of phages [57].

When assessing potential phage candidates for clinical use, 24 hour kinetic growth curves in standard LB at low MOI are routinely used to monitor loss of phage activity following bacterial resistance emergence [58,59]. However, such assays in nutrient-rich laboratory media may not accurately reflect the conditions encountered at the site of infection and this time lapse may be too short to capture resistance emergence, explaining why kinetic growth curves in classical broth are not always predictive for *in vivo* efficacy[58,60]. Our results show distinct patterns between LB and urine with overall higher and more sustained lytic activity observed in urine over 46 hours. This finding suggests that the fitness of the phage-resistant bacteria may be reduced in urine, thus supporting the use of these phages in treatment of UTI. A recent study compared the activity of a phage cocktail against *E. coli* in human urine and a three dimensional urothelial microtissue model [61]. The results showed markedly decreased activity in urine compared to rich-nutrient media, highlighting the importance of infection-site conditions. Interestingly, this study showed that phages modulated host responses without reducing bacterial burden in the microfluidic model, suggesting the importance of phage-immune system interactions [61]. Together, those observations support the need to test phages in conditions as close as possible to the targeted biological environment as a necessary step before *in vivo* application.

Selection of bacterial resistant mutants after exposure to phage is expected *in vivo*, as *in vitro*. However, selection of bacterial mutants with reduced fitness might be a significant biological benefit. Indeed, emerging phage resistance was shown both in clinical use and in pre-clinical data in the urinary environment to be associated with reduced bacterial fitness and antibiotic resensitization [17,62]. This biological trade-off might be of particular interest when treating MDR strains. Preventive as well as curative phage strategies for chronic UTI will need to consider the dynamic interactions between phage, bacteria, and the host defense systems within the bladder. For this *in vivo* modelling is needed to assess activities of the phages in this complex setting. Defining optimal phage dosing, timing, and integration with antibiotics will be essential for ultimately guiding clinical application and advancing phage therapy as a reliable clinical intervention for MDR UTI.

**Table 1.** Genomic and morphological features of EDIRA phages.

|  |  | EDIRA083 | EDIRA088 | EDIRA092 |
| --- | --- | --- | --- | --- |
| Taxonomy |  | yonseivirus | undescribed genus | peekayseptimavirus |
| Capsid | Height (nm) | 55.8±3.6 | 54.9+3.5 | 58.8+1.7 |
|  | Width (nm) | 51.9+3.2 | 53.7+1.8 | 54.2+3.4 |
| Tail | Length (nm) | 189.8+13.3 | 219.1+9.2 | 181.6+2.8 |
|  | Width (nm) | 10.3+0.8 | 17.8+1.5 | 12.9+3.8 |
| Genome size (kb) |  | 57.8 | 48.1 | 50.6 |
| Number of ORFs |  | 88 | 63 | 91 |

## Acknowledgments

We thank the team of the GenoBiomics platform of the AP-HP for performing extraction and sequencing of the microbial genomes and Manuel Majrouh for training and help during electron microscopy acquisition. We are grateful for support for Ultrastructural BioImaging Core Facility equipment from the GIS-IBISA, the DIM One Health, the French government (Agence Nationale de la Recherche) Investissements d’Avenir France BioImaging (FBI, ANR-10-INSB-04-01) and Investissement d’Avenir Laboratoire d’Excellence “Integrative Biology of Emerging Infectious Diseases” (ANR-10-LABX-62-IBEID). This work was partially supported by ANR grant DECOLONIZE ANR-20-AMRB-0004-01 from a FR-DE AMR Bilateral program, ANR grant ANR-22-AAMR-0006 KLEOPATRA under the framework of the JPIAMR - Joint Programming Initiative on Antimicrobial Resistance, Institut Carnot Pasteur Microbes et Santé (ANR 20 CARN 0023-01) and Institut Carnot APHP (ANR 20 CARN 0031-01) and CRC CartoPhage (APHP231314). We thank URC-Est for their support throughout the CRC project. RC research activity was supported through an Interface Contract between Pasteur Institute and Assistance Publique-Hôpitaux de Paris. We are especially grateful to Dr. Molly Ingersoll for constructive support and critical reading of the manuscript.

## Author contributions

RC, CE, and RT conceived and supervised the project, and RC and RT wrote and edited the manuscript. BVB, JG, and CG performed biological experiments; EC, and RT performed electron microscopy; JG, AS, and RT performed genome analysis; GP, CE and LB provided strains and urine; CG, SW, and LM provided water samples for phage isolation; RC, BVB, JG, CE, and RT analyzed data; all authors contributed critical evaluation of the final version of the manuscript.

